# 16S rRNA gene sequences of *Candidatus* Methylumidiphilus (*Methylococcales*), a putative methanotrophic genus in lakes and ponds

**DOI:** 10.1101/2021.09.28.462094

**Authors:** Antti J Rissanen, Moritz Buck, Sari Peura

## Abstract

A putative novel methanotrophic genus, *Candidatus* Methylumidiphilus (*Methylococcales*), was recently shown to be ubiquitous and one of the most abundant methanotrophic genera in water columns of oxygen-stratified lakes and ponds of boreal and subarctic area. However, it has probably escaped detection in many previous studies using 16S rRNA gene amplicon sequencing due to insufficient database coverage, which is because *Ca*. Methylumidiphilus lacks cultured representatives and previously analysed metagenome assembled genomes (MAGs) affiliated with it do not contain 16S rRNA genes. Therefore, we screened MAGs affiliated with the genus for their 16S rRNA gene sequences in a recently published lake and pond MAG dataset. Among 66 MAGs classified as *Ca*. Methylumidiphilus (with completeness over 40% and contamination less than 5%) originating from lakes in Finland, Sweden and Switzerland as well as from ponds in Canada, we could find 5 MAGs each containing one 1532 bp long sequence spanning the V1-V9 regions of the 16S rRNA gene. After removal of sequence redundancy, this resulted in two unique 16S rRNA gene sequences. These sequences represented two different putative species, i.e. *Ca.* Methylumidiphilus alinenensis (Genbank accession: OK236221) as well as another so far unnamed species of *Ca*. Methylumidiphilus (Genbank accession: OK236220). We suggest that including these two sequences in reference databases will enhance 16S rRNA gene - based detection of members of this genus from environmental samples.

## 1. INTRODUCTION

Methanotrophic bacteria are widely distributed and play a crucial role in consuming greenhouse gas methane in natural (wetlands, lakes, oceans, soils) and anthropogenic (wastewater treatment plants, landfills) methane-producing ecosystems (Hanson & Hanson 1996, Kallistova et al. 2005). Currently, their identity, diversity and community structure are mostly studied using polymerase chain reaction (PCR)-based techniques, i.e. high-throughput amplicon sequencing and quantitative PCR, targeting the 16S rRNA gene or the *pmoA* gene encoding the beta subunit of particulate methane monooxygenase (Rissanen et al. 2018, Mayr et al. 2020a, b). The advantage of these PCR-based methods is their cost-effectiveness and speed in the analyses of multiple samples. Yet, recently, more expensive, shotgun metagenomic study methods, which overcome the problem of primer bias/mismatch inherent in PCR-based methods and which allow also insights into the genetic potential of the *in situ* bacteria, have been employed in studies of methanotrophic communities (Rissanen et al. 2018, Smith & Wrighton 2019, van Grinsven et al. 2020).

The outcome of DNA sequencing – based taxonomic analyses are dependent on the quality and taxonomic coverage of reference database(s). Using PCR-free, 16S rRNA gene - independent shotgun metagenomic sequencing approach, we recently showed that a putative novel genus of methanotrophs, *Candidatus* Methylumidiphilus (order *Methylococcales*), was ubiquitous and one of the most abundant methanotrophic genera in water columns of oxygen-stratified lakes and ponds of boreal and subarctic area (Rissanen et al. 2018, 2020, Martin et al. 2021). The first putative species of this genus was named as *Candidatus* Methyloumidiphilus alinensis [the name later proposed to be changed to *Ca*. Methylumidiphilus alinenensis (Oren et al. 2020)], which was represented by an abundant metagenome assembled genome (MAG) in the water samples of boreal Lake Alinen Mustajärvi (Rissanen et al. 2018). Interestingly, depending on the reference database, our analyses suggested that the genus had probably not been classified as a methanotroph (*Methylococcales*) at all (i.e. classified as unclassified *Gammaproteobacteria*), or had been classified only at the order level (i.e. as unclassified *Methylococcales*) in previous 16S rRNA gene - based studies (Rissanen et al. 2018, Martin et al. 2021). To aid in correctly classifying the 16S rRNA genes of this genus, a previously published clone library sequence from L. Alinen Mustajärvi (Genbank, HE616416, 830bp) was determined to represent *Ca*. Methylumidiphilus alinenensis by Rissanen et al. (2018), and was used as a database sequence in some subsequent 16S rRNA gene analyses (Thamdrup et al. 2019, Rissanen et al. 2020). However, the 16S rRNA gene - based phylogenetic position of *Ca*. Methylumidiphilus remains to be confirmed (Knief 2019), as 16S rRNA gene sequences are not available from the previously reconstructed MAGs representing the genus (Rissanen et al. 2018, 2020). In addition, HE616416 covers only V1-V5 regions of the 16S rRNA gene making it impossible to use it as a reference sequence in studies focusing on V6-V9 regions. Modern PCR-based amplicon sequencing analyses using long-read sequencing technologies (PacBio or Oxford Nanopore) covering the whole V1-V9 regions of 16S rRNA gene as well as PCR-free shotgun metagenomic – based 16S rRNA gene analyses would also require full-length or almost full length 16S rRNA gene sequences as references.

Metagenomic assembly and binning approaches often fail in reconstruction of the 16S rRNA genes of the target organisms, for example lake methanotrophs (Rissanen et al. 2020), yet in some rare cases it has been successful (van Grinsven et al. 2020). Therefore, screening of multiple MAGs representing the organism(s) of interest might be needed to identify MAGs containing reconstructed 16S rRNA genes. Our previously mentioned observation on the ubiquitousness and abundance of *Ca*. Methylumidiphilus was based on analyses with the recently published shotgun metagenomic dataset from water columns of lakes and ponds (Buck et al. 2021, Martin et al. 2021). Therefore, with the aim to provide 16S rRNA gene sequences representing *Ca*. Methylumidiphilus to be included in reference databases, we screened the MAGs affiliated with *Ca*. Methylumidiphilus in this dataset for their 16S rRNA genes.

## 2. MATERIALS AND METHODS

We used previously published MAG dataset from 41 stratified lakes and ponds mainly located in the boreal and subarctic regions, but also from one tropical reservoir and one temperate lake (Buck et al. 2021). See Buck et al. (2021) on detailed report of the sample collection, DNA extraction, library preparation, sequencing and bioinformatic analyses (trimming/filtering, assembly, metagenomic binning). Furthermore, Buck et al. (2021) used checkM (v. 1.0.13) for assessing the prokaryotic completeness and redundancy of the MAGs (Parks et al. 2015), while GTDB-Tk (version 102 with database release 89) (Parks et al. 2018) as well as SourMASH’s lca classifier (Brown & Irber 2016) were used for their taxonomic classification. Finally, Buck et al. (2021) clustered the MAGs, starting with 40% complete genomes with less than 5% contamination, into metagenomic operational taxonomic units (mOTUs) at 95 % level of average nucleotide identity (ANI) calculated using fastANI (v. 1.3) (Jain et al. 2018).

For our analyses, we chose MAGs with genus level taxonomic classification of “d__Bacteria;p__Proteobacteria;c__Gammaproteobacteria;o__Methylococcales;f__Methylococcaceae;g__AMB10-2013” and which had completeness over 40% and contamination less than 5%. The genus level name “g__AMB10-2013” denotes the MAG of *Candidatus* Methylumidiphilus alinenensis (GCA_003242955) discovered from water column of boreal lake Alinen Mustajärvi (Rissanen et al. 2018). The chosen MAGs were functionally annotated using Prokka (v. 1.14.6) (Seemann 2014), which included detection of rRNA genes using barrnap (v. 0.9) (Seemann 2018). Phylogenetic trees based on 16S rRNA genes were built using the maximum likelihood algorithm (generalized time reversible model) with 100 bootstraps in Mega × (Kumar et al. 2018). Furthermore, phylogenomic trees including reference genomes as well as representative MAGs of mOTUs affiliated to *Ca*. Methylumidiphilus (i.e. with genus level taxonomic classification of “g__AMB10-2013”) were built using PhyloPhlAn (version 3.0.58) with the PhyloPhlAn database (400 universal marker genes) (Asnicar et al. 2020).

## 3. RESULTS AND DISCUSSION

In Buck et al. (2021) dataset, there were 66 MAGs, which had completeness over 40% and contamination less than 5% and with taxonomic assignment to *Ca.* Methylumidiphilus (i.e. “g__AMB10-2013”). These MAGs were classified into 12 mOTUs (Fig. 1), whose representative genomes originated from lakes in Finland, Sweden and Switzerland as well as from ponds in Canada, further reflecting the ubiquitousness and wide geographical distribution of the genus (Buck et al. 2021, Martin et al. 2021). Of the mOTUs, mOTU 0341, 2711, 1471, 1599 and 2021 were represented by more than one MAG, i.e. 42, 6, 4, 4 and 3 MAGs, respectively, while each of the other mOTUs included only one MAG (Fig. 1). Our previously studied MAGs of *Ca*. Methylumidiphilus originating from boreal lakes, i.e. *Ca*. Methylumidiphilus alinenensis from Lake Alinen Mustajärvi and bin-0959 from L. Lovojärvi, were also included in phylogenomic tree analysis, with a result indicating that they belong to mOTUs 0341 and 1599, respectively (Fig. 1) (Rissanen et al. 2018, 2020).

**Fig. 1.**
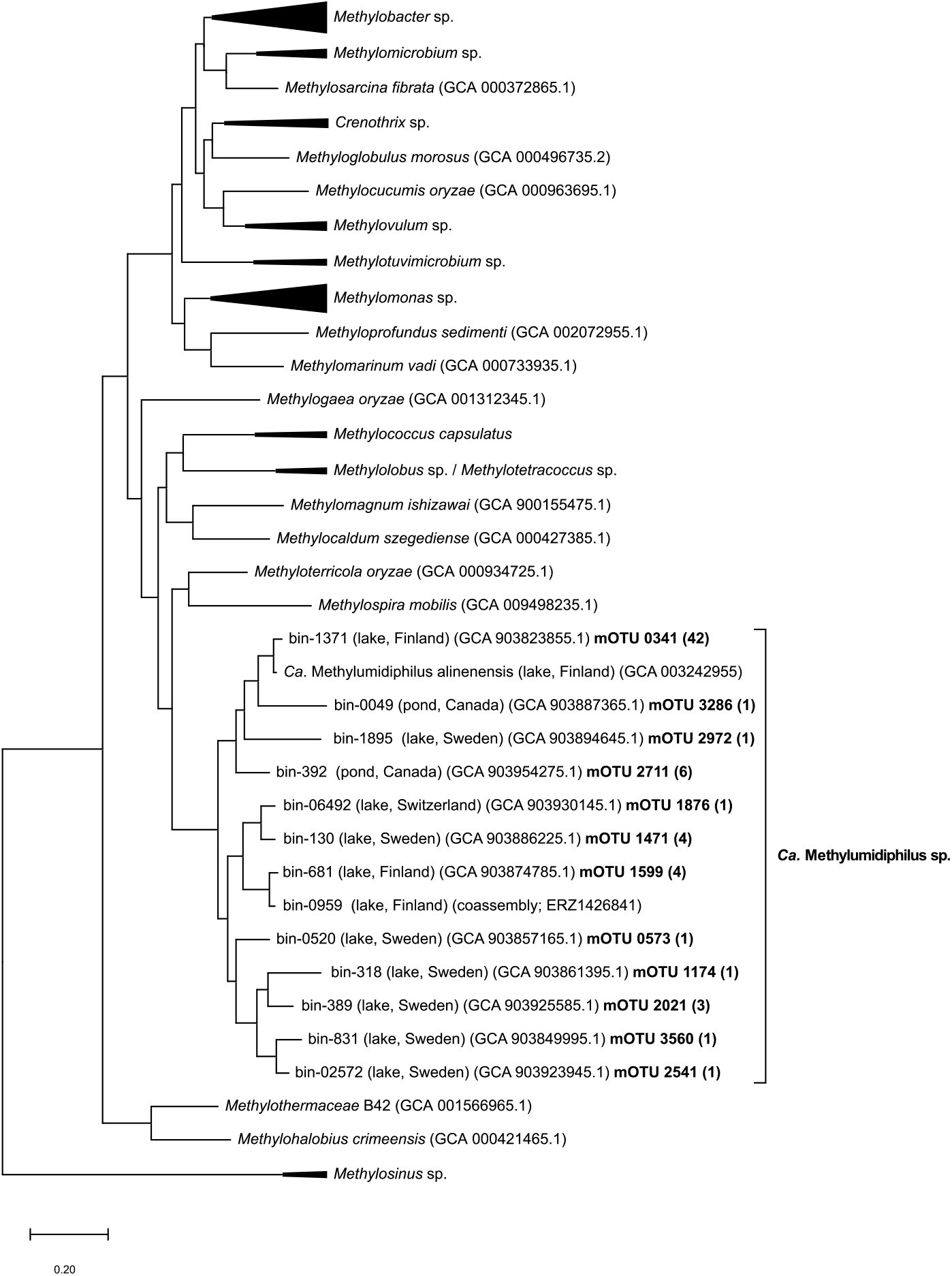
Phylogenomic tree (PhyloPhlAn) of *Methylococcales* (outgroup = alphaproteobacterial genus *Methylosinus*). The tree shows representative metagenome assembled genomes (MAGs) of metagenomic operational taxonomic units (mOTU) affiliated with *Candidatus* Methylumidiphilus in Buck et al. (2021) dataset, as well as MAGs, which we analysed previously, i.e. *Ca*. Methylumidiphilus alinenensis and bin-0959 (Rissanen et al. 2018, 2020). mOTU number as well as the number of MAGs belonging each of the mOTU (in brackets after mOTU number) are highlighted with bold text

Fragments of 16S rRNA genes were found within 15 out of the studied 66 MAGs. Of these, 6 MAGs included almost full length 16S rRNA gene sequences (1530-1532 bp) and were chosen for further analyses, while all other had length less than 1200 bp. In the preliminary taxonomic classification analyses using blastn (Altschul et al. 1990), one of the 16S rRNA gene sequences (from bin-1515 GCA_903920655.1) was closely affiliated with *Methylobacter* (98.5 % identity with *Methylobacter tundripaludum* SV96, NR_042107), and hence probably came from a wrongly binned contig, while the other 5 had high identity (96.0 – 99.6 % identity) with the partial 16S rRNA gene sequence HE616416 previously suggested to represent *Ca*. Methylumidiphilus alinenensis (Rissanen et al. 2018). The phylogenetic tree based on 16S rRNA gene sequences confirmed the phylogenetic position of these sequences as they formed a distinct cluster, with *Methyloterricola* and *Methylospira* as their neighbouring genera (Fig. 2), which agrees with previous phylogenetic analyses with HE616416 (Rissanen et al. 2018, Knief 2019). The 16S rRNA gene sequences formed two clusters, one including three identical 16S rRNA gene sequences representing mOTU 0341 (submitted to Genbank with accession: OK236221), and the other including two identical 16S rRNA gene sequences representing mOTU 2711 (Genbank accession: OK236220) (Fig. 2). The blastn-analysed identities of the 16S rRNA gene sequences of these clusters to those of *Methylospira palustris* (90.9% and 90.8% identity for mOTUs 2711 and 0341, respectively) and *Methyloterricola oryzae* (91.1% and 91.6% identity for mOTUs 2711 and 0341, respectively) were much lower, while their identities to each other (97.5% identity between mOTU 2711 and 0341) were much higher, than the suggested 94.5% identity threshold to delineate different genera (Yarza et al. 2014), further confirming their taxonomic assignment to a different genus than *Methylospira* and *Methyloterricola*. Phylogenomic analyses as well as the high identity of the 16S rRNA gene sequences of mOTU 0341 to HE616416 (99.6% identity) further suggests that mOTU 0341 represents *Ca*. Methylumidiphilus alinenensis (Fig. 1). In addition, both phylogenomic as well as 16S rRNA gene analyses suggest that mOTU 2711 represents a different, so far unnamed, species of *Ca*. Methylumidiphilus.

**Fig. 2.**
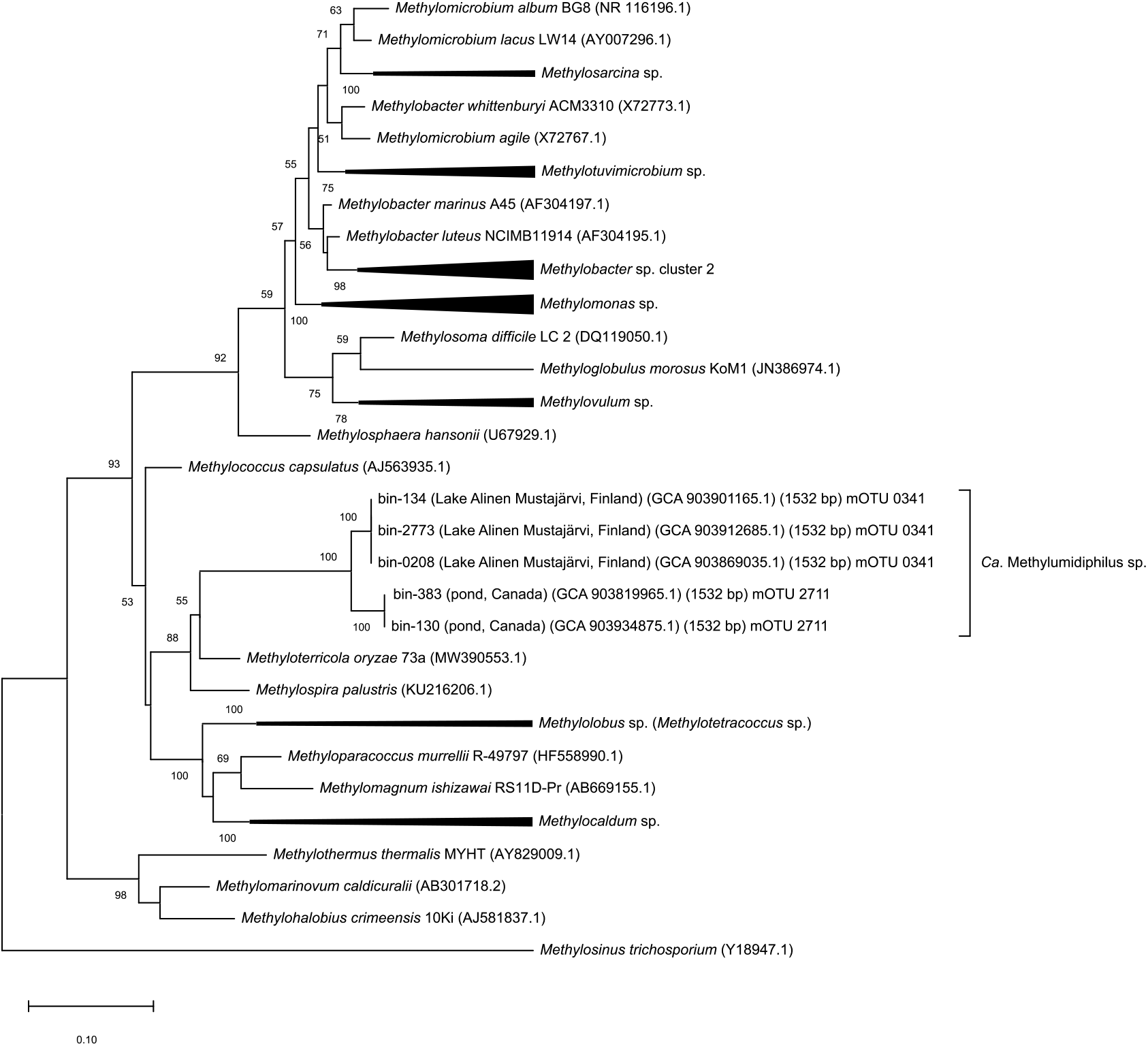
Phylogenetic tree based on 16S rRNA genes of *Methylococcales* (outgroup = alphaproteobacterial genus *Methylosinus*). The three shows 16S rRNA gene sequences, spanning V1-V9 regions of the 16S rRNA gene, detected in 5 MAGs affiliated with *Ca*. Methylumidiphilus. The mOTU number of the MAGs is also shown (see Fig. 1)

In this study, we provided for the first time almost full length 16S rRNA gene sequences representing the putative methanotrophic genus, *Ca*. Methylumidiphilus, which is ubiquitous in lakes and ponds of boreal and subarctic area (Martin et al. 2021), and according to this study, is also present in a temperate lake, Lake Loclat, in Switzerland (Fig. 1). MAG of *Ca*. Methylumidiphilus alinenensis is also the first representative genome of the Lake Washington (LW) – cluster, which is detected in *pmoA* – gene based phylogenetic analyses, and which lacks cultured representatives (Knief 2019, Rissanen et al. 2020). LW cluster generally includes *pmoA* sequences from different aquatic habitats (Knief 2019). Hence, we suggest that including the provided 16S rRNA gene sequences in reference databases will enhance the 16S rRNA gene – based detection of members of *Ca*. Methylumidiphilus in further studies of methanotrophic communities of lakes and other aquatic ecosystems.

## Acknowledgements

This study was supported by Kone Foundation (Grant No. 201803224) for AJR. The sequencing was funded by a grant from the Science for Life Laboratory biodiversity program and SciLifeLab fellows program. The computations were performed on resources provided by SNIC through Uppsala Multidisciplinary Center for Advanced Computational Science (UPPMAX) under Project SNIC snic2020-5-19.

